# Freshwater ‘microcroissants’ shed light on a novel higher-level clade within Trebouxiophyceae and reveal the genus *Chlorolobion* as a trebouxiophyte

**DOI:** 10.1101/2024.01.20.576396

**Authors:** Dovilė Barcytė, Ladislav Hodač, Marek Eliáš

## Abstract

Trebouxiophyceae is a widespread and species-rich green algal class encompassing mostly coccoid algae with a simple ovoid or ellipsoidal outline. However, some poorly-sampled lineages have evolved more elaborate shapes or even complex thalli, adding to the class’s morphological diversity. Led by new and previously established strains, this study additionally uncovered a clade of croissant-like trebouxiophytes. Phylogenetic analyses inferred from nuclear 18S rDNA and chloroplast *rbcL* sequences confirmed the monophyly of the ‘microcroissant’ clade, which we propose to be classified as a new family, Ragelichloridaceae. This family includes two novel genera, *Ragelichloris* and *Navichloris*, and the previously described *Thorsmoerkia*. The position of Ragelichloridaceae within Trebouxiophyceae stayed unresolved but chloroplast phylogenomics showed that the family belongs to the broader *incertae sedis* group, which also includes *Xylochloris* and *Leptosira*. In addition, our study showed that the microcroissant-like genus *Chlorolobion*, previously classified within Chlorophyceae, is a genuine trebouxiophyte, potentially related to Ragelichloridaceae.

**Highlights:**

- A new family-level clade uncovered within Trebouxiophyceae.
- Two new genera described.
- The genus *Chlorolobion* shown to be a trebouxiophyte.

## Introduction

Trebouxiophyceae is one of the major groups of green algae and encompasses organisms with a broad range of lifestyles – from free living photoautotrophs and endosymbiotic partners to heterotrophic parasites. They also exhibit a great diversity in morphological features, though the most widespread morphotype is a simple coccoid form. In fact, cells with a spherical, ovoid, or ellipsoidal outline form a vast majority of the group’s described species. This morphology appears to represent the plesiomorphic state, as it occurs across the trebouxiophyte phylogenetic tree, intermingled with lineages exhibiting different, more intricate morphotypes (Leliaert *et al*., 2012). Therefore, morphological traits alone are highly unreliable in inferring phylogenetic relationships or defining higher-order taxa in Trebouxiophyceae. A great example of contrasting morphologies is the order Microthamniales encompassing both the multicellular branched filamentous *Microthamnion* and the unicellular coccoid *Coleochlamys* revealed as sister taxa only thanks to molecular phylogenetics (Lemieux *et al*., 2014; Barcytė *et al*., 2021). Another similar example is the *Prasiola* clade accommodating organisms with a wide range of thallus morphologies and organization, spanning from unicellular through filamentous to pseudoparenchymatous (Pröschold & Darienko, 2020). On the other hand, the order Watanabeales, first recognized as a phylogenetic clade of *Chlorella*-like representatives (i.e. *Watanabea* clade; Karsten *et al*., 2005), was subsequently solidified as a formal taxon when reproduction strategies among close relatives were compared (Li *et al*., 2021).

Despite the growing knowledge of trebouxiophyte diversity, with new and previously overlooked taxa (species and genera) being described at a steady pace (e.g., Barcytė *et al*., 2017; Li *et al*., 2020; Malavasi *et al*., 2022; Kato *et al*., 2023), the higher-level phylogenetic relationships within Trebouxiophyceae, and formal delimitation of taxa at the suprageneric level, remain highly unsettled. Identifying monophyletic clades with a full statistical support is primarily hampered by unstable phylogenetic relationships largely caused by poorly sampled lineages “jumping” across the Trebouxiophyceae phylogenies depending on the taxa sampled or a molecular marker used. Phylogenomics could overcome this problem and generate a reliable backbone phylogeny as well as shake or reshape our understanding of some of the major groupings (Lemieux *et al*., 2014). However, for many understudied lineages multigene data are so far missing and nuclear 18S rDNA trees (sometimes combined with phylogenies derived from the chloroplast *rbcL* gene) remain the primary means to infer the relationships within the group.

Since “green balls” represent most of the Trebouxiophyceae diversity, organisms with conspicuous morphology attract attention in terms of their phylogenetic placement within the class. In some cases, such organisms form deeply branching isolated lineages. A notable example is the recently described semi-aerophytic crescent-shaped *Thorsmoerkia curvula*, which was not assigned to any of the existing orders or clades and allegedly lacked sequenced close relatives (Nicoletti *et al*., 2021). Another example of an enigmatic and phylogenetically unresolved lineage is represented by a *Navicula*-shaped coccoid alga (strain SAG 2477) isolated from soil in Germany, provisionally considered a candidate new species in a new genus in a PhD thesis (Hodač, 2015). This alga is phylogenetically close to the organism hiding under the name BCP-BC4VF9, isolated and sequenced from the Central Desert in Baja California, Mexico (Fučíková *et al*., 2014). Unfortunately, the BCP-BC4VF9 strain did not survive and its fine morphological features stayed undocumented. Finding sister taxa of such orphan trebouxiophyte lineages could not only help to place them better among other subgroups but also assist in stabilizing the whole phylogenetic backbone of the class, a prerequisite for a better understanding of the Trebouxiophyceae evolution.

We recently isolated two peculiar *Sphagnum*-associated croissant-shaped algal strains that we aimed to place in the green algal phylogeny and uncover their closest relatives. Utilizing 18S rDNA and *rbcL* as phylogenetic markers, we found out these new isolates to belong to Treboxiophyceae, specifically to a broader novel trebouxiophyte clade additionally including the aforementioned enigmatic lineages. Microscopic observations confirmed a similar morphology of the three distinct genus-level lineages (*Thorsmoerkia*, *Ragelichloris* gen. nov., and *Navichloris* gen. nov.) constituting the novel clade, suggesting the croissant-shaped trebouxiophytes be recognized as a novel higher-level taxon within the class. This notion was further supported by the multigene phylogenomic analysis based on the chloroplast genome data, demonstrating the deeply branching nature of the croissant lineage within Trebouxiophyceae, forming sister relationships with other *incertae sedis* lineages of the class. Finally, sequence analysis of the similar morphotype-bearing genus *Chlorolobion* (including the type species *C. obtusum*) provided the first evidence of its affiliation to Trebouxiophyceae and elucidated its polyphyletic nature, with two of the species belonging to the novel ‘microcroissant’ clade.

## Materials and methods

### Strains and microscopy

Strain “PLY” was isolated from a *Sphagnum* moss sample taken in October 2021 from the raised bog Plynoja (55°19′23″ N, 22°8′18″ E) situated in the Pagramantis Regional Park, Tauragė, Lithuania (Fig. S1). Strain “VMJ” was isolated from a *Sphagnum* sample taken in May 2022 from the shore of the Great Moss Lake (Velké mechové jezírko; 50°13′10.92″ N, 17°17′12.84″ E) situated in the National Natural Reserve Rejvíz, Czech Republic (Fig. S2). Based on a preliminary phylogenetic analysis, we additionally obtained and studied strain SAG 2477 (Sammlung von Algenkulturen Göttingen), deposited to the collection as an “unidentified Trebouxiophyte”. Motivated by morphological identification of one of the new isolates, we also purchased nine ACOI (Coimbra Collection of Algae) strains identified by the curator as five different species of the genus *Chlorolobion* (Table 1). All strains were cultivated in liquid Bold’s Basal Medium (BBM; Bischoff & Bold Citation 1963) in 25 cm^2^ cell culture flasks with filter caps (SPL Life Sciences), as well as on agarised BBM medium (1.5% agar) in Petri dishes.

**Table 1.** SAG and ACOI strains evaluated for their connection to the ‘microcroissant’ clade.

| Strain | Original identification | Habitat, collection site | Updated species name | Classification |
| --- | --- | --- | --- | --- |
| SAG 2477 | unidentified Trebouxiphyte | soil in spruce forest, Swabian Alb, Germany | *Navichloris terrestris* | Ragelichloridaceae, Trebouxiphyceae |
| ACOI 732 | *Chlorolobion obtusum* | Serra da Estrela, Covão do Boeiro, Portugal |  | Trebouxiphyceae, *incertae sedis* |
| ACOI 231 | *Chlorolobion lunulatum* | pond near the river, São João do Campo, Portugal |  | Selenastraceae, Chlorophyceae |
| ACOI 811 | *Chlorolobion lunulatum* | puddle, Serra do Gerês, Leonte, Portugal |  | Selenastraceae, Chlorophyceae |
| ACOI 2681 | *Chlorolobion lunulatum* | puddle, Serra do Gerês, Portugal | *Thorsmoerkia lunulata* | Ragelichloridaceae, Trebouxiphyceae |
| ACOI 3192 | *Chlorolobion lunulatum* | puddle, Serra do Gerês, Leonte, Portugal | *Thorsmoerkia lunulata* | Ragelichloridaceae, Trebouxiphyceae |
| ACOI 208 | *Chlorolobion saxatile* | pond near Arazede, Amieiro, Portugal |  | Selenastraceae, Chlorophyceae |
| ACOI 3295 | *Chlorolobion guanense* | unknown |  | Selenastraceae, Chlorophyceae |
| ACOI 820 | *Chlorolobion braunii* | Leiria, Portugal |  | Selenastraceae, Chlorophyceae |
| ACOI 2567 | *Chlorolobion braunii* | pond, Castelo Branco, Penha Garcia, Portugal |  | Selenastraceae, Chlorophyceae |

Light microscopy was carried out with an Olympus BX53 light microscope (Tokyo, Japan). Micrographs were taken with an Olympus DP73 digital camera (Tokyo, Japan). The Olympus cellSens Imaging Software v1.6 was used to process images and obtain morphometric measurements of the cells. In addition, strain PLY was further investigated using transmission electron microscopy (TEM). For TEM, a flat specimen carrier of 3 mm diameter (Leica Microsystems, Vienna, Austria) was first filled with a drop of 20% BSA (bovine serum albumin) cryoprotectant, followed by a drop of the centrifuged cell suspension. The specimen carriers were then rapidly frozen (within 30 msec at a pressure of 2,100 Pa) using a Leica EM ICE High Pressure Freezer (Leica Microsystems). Following freezing, the flat specimen carriers were transferred to screw-capped containers filled with a pre-cooled (-90°C) substitution medium (2% osmium tetroxide in 100% acetone). The containers were placed into a freeze substitution device Leica AFS (Leica Microsystems) pre-cooled to -90°C. The samples were substituted for a week started at - 90°C for 96 h, followed by warming at 5°C/h to -20°C. After 24 h at -20°C, warming was continued at 3°C/h to 3°C. This was followed by 8 h at 3°C and 18 h at 4°C. After a week the specimen carriers were rinsed three times with 100% acetone at room temperature. During the rinsing procedure round pieces of cell suspensions were released from the specimen carriers and those pieces of cells were infiltrated into the resin. Finally, ultrathin sections of the resin samples were cut on a Reichert-Jung Ultra-cut E ultramicrotome (Wien, Austria) and stained using uranyl acetate and lead citrate. Sections were examined using a JEOL JEM-1011 electron microscope (Tokyo, Japan).

### DNA extraction, PCR, and sequencing

Total genomic DNA from each strain was extracted using a Geneaid Plant Genomic DNA Mini Kit (New Taipei City, Taiwan). Nuclear 18S rDNA sequences were PCR-amplified with the primer pairs 18SF plus 18SR (Katana *et al*., 2001) or 20F (Thüs *et al*., 2011) plus CH1750R (Hallmann *et al*., 2013). A segment of the chloroplast RuBisCO large subunit gene (*rbcL*) was amplified with primers M35F plus M1338R (McManus & Lewis, 2011) and rbcL1F plus rbcL23R (Hoham *et al*., 2002). The internal transcribed spacer 2 (ITS2) rDNA region was amplified using the primers AL1500af (Helms *et al*., 2001) plus LR3 (Vilgalys & Hester, 1990). All PCR reaction mixtures were prepared using either 2x PCRBIO Taq Mix Red (PCR Biosystems, London, UK) or Qiagen Multiplex PCR Plus Kit (Hilden, Germany). The PCR amplification thermal profile for each primer pair is listed in Table S1. The PCR products were purified with QIAquick PCR & Gel Cleanup Kit (Qiagen, Hilden, Germany). All amplicons were directly Sanger-sequenced at Eurofins Genomics (Ebersberg, Germany). The primer pair mci01_F (5′ ACAAATCACAAGCAGAAACGGG 3′) plus mci01_R (5′ AATGTCCCTTAACCTCCAAATAAGG 3′) were additionally designed for sequencing the *rbcL* intron. The obtained reads were assembled using SeqAssem ver. 07/2008 (Hepperle, 2017). Sequences were deposited in GenBank and are available under accession numbers XX000000–XX000000.

### Phylogenetic analyses

In addition to newly acquired sequences, this study compiled 18S rDNA and *rbcL* sequences from GenBank (www.ncbi.nlm.nih.gov/genbank/) to represent all major known lineages of the class Trebouxiophyceae, with Chlorophyceae representatives serving as outgroup taxa. The 18S rDNA dataset was aligned with the online MAFFT version 7 (https://mafft.cbrc.jp/alignment/server/) using the L-INS-i method (Katoh & Standley, 2013). The resulting alignment was trimmed with trimAl v1.2rev59 using the -automated1 mode (Capella-Gutiérrez *et al*., 2009), yielding the final dataset of 83 sequences with 1,686 aligned positions. Maximum likelihood (ML) analysis was performed with IQ-TREE multicore version 2.2.5 (Nguyen *et al*., 2015) using the TIM2e+I+R4 model and a non-parametric bootstrapping with 100 replicates. The evolutionary model was automatically selected by ModelFinder (Kalyaanamoorthy *et al*., 2017) implemented in IQ-TREE. Bayesian inference (BI) was carried out with MrBayes v3.2.7 (Ronquist *et al*., 2012) using the GTR+I+G model and two independent runs with one cold and three heated chains each ran for 3,000,000 generations and sampled every 100th generation. The first 25% sampled datapoints were discarded as burn-in. Parameter stability and run convergence were assessed using Tracer v1.7.2 (Rambaut *et al*., 2018). The *rbcL* dataset of 73 sequences was aligned using the same method, resulting in 1,431 aligned positions. For the ML analysis, this dataset was partitioned to the 1st, 2nd, and 3rd codon positions, and respective models GTR+F+R4, SYM+I+R2, and TIM3+F+I+R4 were applied to each partition. Non-parametric bootstrapping technique with 100 replicates was used to assess node support. For BI analysis, GTR+I+G model was set to each codon position, followed by the steps described above.

For phylogenomic analysis, we extracted plastome-derived transcript sequences from a transcriptome assembly generated from strain PLY (to be published elsewhere) and deduced the corresponding encoded protein sequences. These were included into 79 chloroplast genome-encoded protein datasets assembled previously by Turmel et al. (2016) (see also Table S2). The same datasets were further updated by including *Kalinella pachyderma*, *Massjukichlorella minus*, and *Medakamo hakoo* (Takusagawa *et al*., 2021; Liu *et al*., 2023). Single-gene alignments were prepared using the strategy outlined above and were subsequently concatenated into a single dataset using FASconCAT-G_v1.05 (Kück & Longo, 2014). ML analysis was carried out employing the automatically selected Q.yeast+I+G4 model and a non-parametric bootstrapping with 100 pseudoreplicates.

### ITS2 rDNA secondary structures

The ITS2 region was annotated using the ITS2 database (Merget *et al*., 2012). The secondary structure of each sequence was generated using the UNAFold web server (Zuker, 2003), and the minimum energy structure was exported as a Vienna-formatted text file. The structures were inspected and edited in 4SALE v1.7.1 (Seibel *et al*., 2006) to correct misfolded regions. Sequences with their secondary structures were aligned using the same program and the ClustalW algorithm. The resulting alignment was used to compute a matrix of compensatory base changes between all pairs of sequences and structures (in 4SALE), and to calculate pairwise phylogenetic distances in the program ProfDist v0.9.9. (Wolf *et al*., 2008). The retrieved phylogenetic distances between sequences + structures were graphically visualised as a UPGMA cladogram without constraints (cophenetic correlation = 0.53) in the program PAST 4.14 (Hammer *et al*., 2001). Folded ITS2 structures were visualised using the online tool Pseudoviewer (http://pseudoviewer.inha.ac.kr) with default settings. Graphical elements were adapted in Inkscape 1.1.

## Results

### Morphology

Vegetative cells of strain PLY were crescent-shaped with narrowly rounded ends (Figs 1, 2). In mature cells, the ends we also slightly bent. Some cells exhibited a tapered shape, being wider at one end than the other (Figs 2, 3). Cell wall was smooth and robust. The cells were 12–20 µm long and 3–8 µm wide. They contained a single chloroplast without a clearly seen pyrenoid. In young cells the chloroplast was trough-shaped and occupied a half of the cell volume. With age, the chloroplast became lobed into two or more parts (Figs 1–3). Small starch grains were seen squeezed between thylakoids under TEM (Fig. 2). The nucleus was on the opposite side to the chloroplast, in the central part of the cell. Oil droplets were also observed in some cells (Fig. 3). When cultivated on agar plates, a majority of cells became drop-like with one end prominently broader than the other (Fig. 4). Reproduction took place by autosporulation, forming two, four or eight daughter cells within a single mother cell. The daughter cells were released by a rupture of the mother cell wall (Fig. 5).

**Fig. 1.**
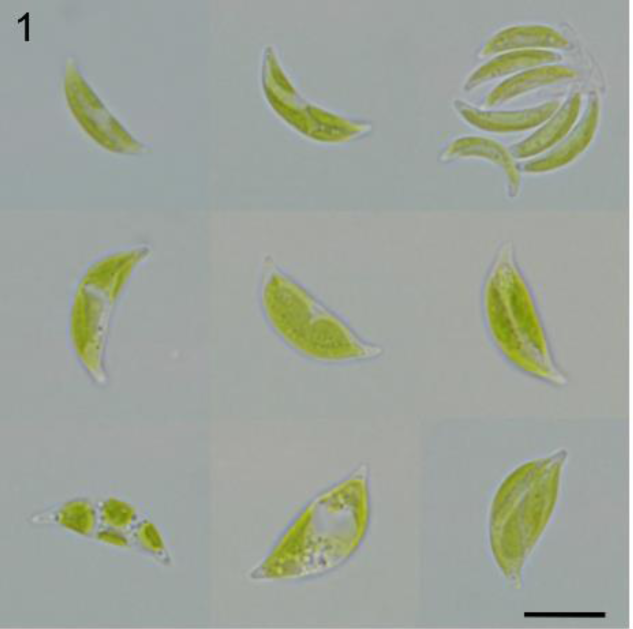
Morphology and ultrastructure of the newly isolated strain PLY (*Ragelichloris palustris* gen. et sp. nov.). Light micrographs of the strain cultivated in liquid medium.

**Fig. 2.**
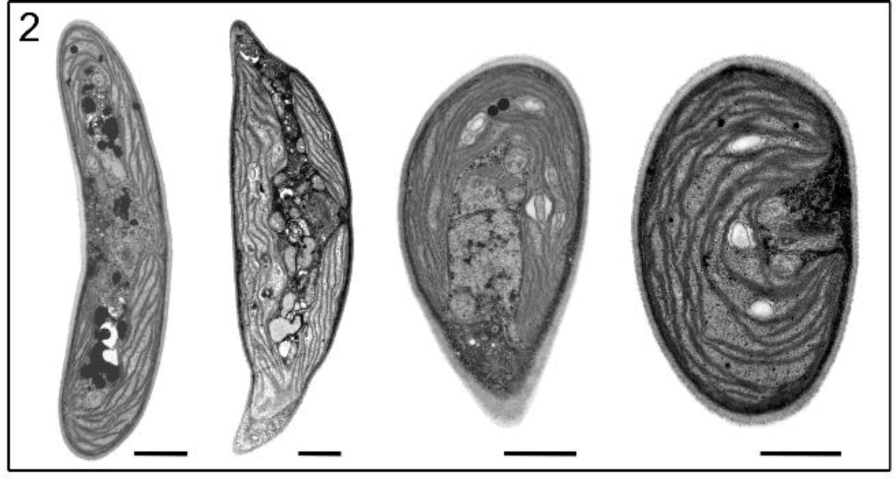
Morphology and ultrastructure of the newly isolated strain PLY (*Ragelichloris palustris* gen. et sp. nov.). Ultrastructure of vegetative cells.

**Fig. 3.**
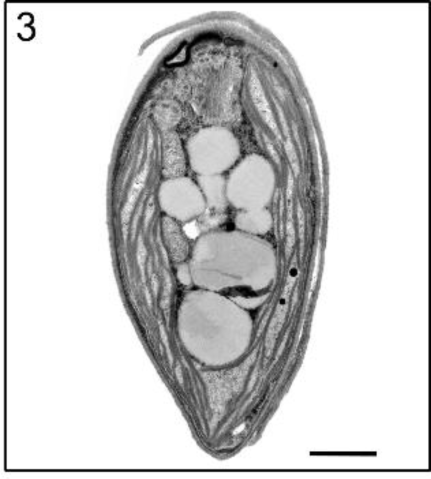
Morphology and ultrastructure of the newly isolated strain PLY (*Ragelichloris palustris* gen. et sp. nov.). Vegetative cell full of lipid droplets.

**Fig. 4.**
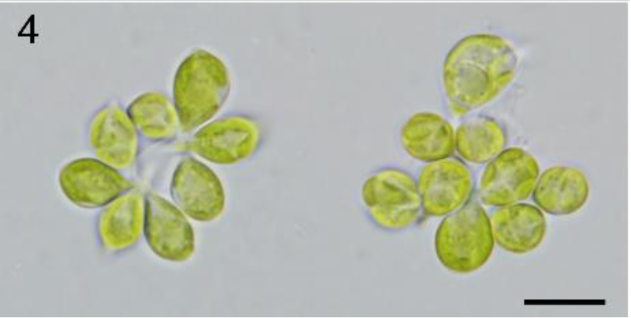
Morphology and ultrastructure of the newly isolated strain PLY (*Ragelichloris palustris* gen. et sp. nov.). Cells grown on agar plates.

**Fig. 5.**
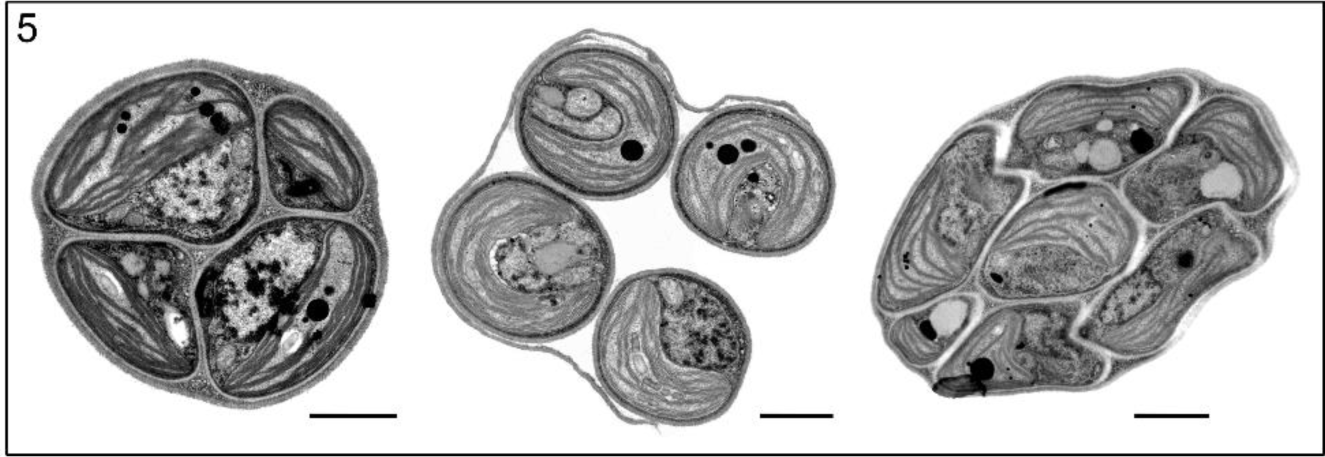
Morphology and ultrastructure of the newly isolated strain PLY (*Ragelichloris palustris* gen. et sp. nov.). Autosporangia with four and eight daughter cells. Scale bars = 10 µm in Figs 1, 4 and 1 µm in Figs 2, 3, 5.

Vegetative cells of strain VMJ were also crescent-shaped with narrowly rounded ends (Figs 6, 7). One cell end was often slightly slenderer than the other. Their size ranged from 6–24 µm in length and 3–8 µm in width. A trough-shaped chloroplast occupied most of the cell volume. The pyrenoid was often obscure in liquid-grown culture of the strain (Fig. 6). However, single pyrenoids became well discernible in cells grown on agar plates (Fig. 7; arrowheads). Cells were frequently vacuolated, with either two prominent vacuoles or a chain of smaller ones (Fig. 7). A single nucleus was present within the cell. The reproduction occurred by producing four to eight autospores per autosporangium.

**Fig. 6.**
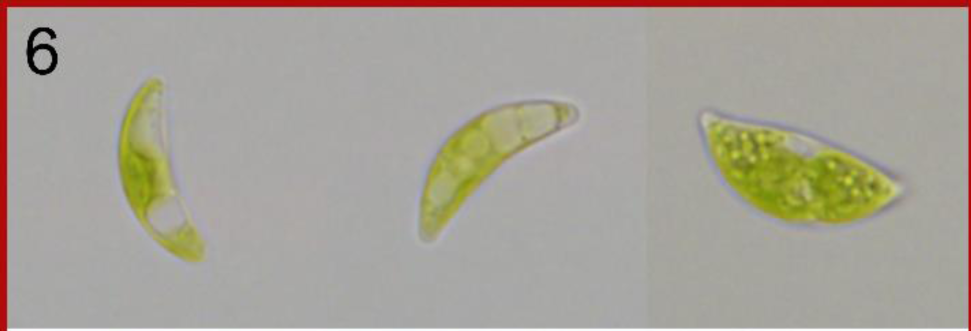
Morphology of the ‘microcroissant’ clade. VMJ strain in liquid medium.

**Fig. 7.**
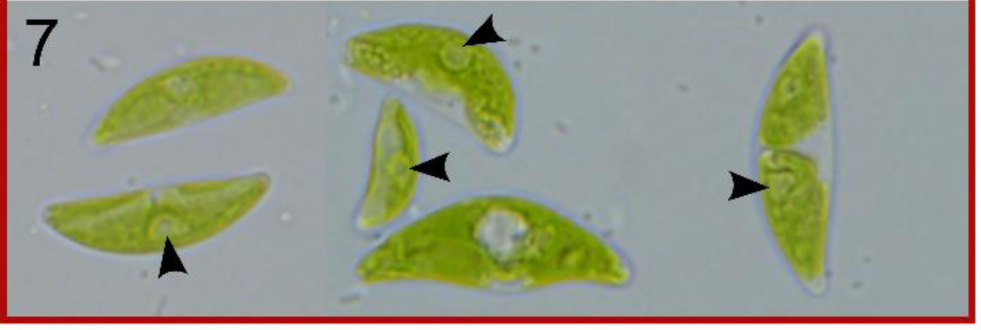
Morphology of the ‘microcroissant’ clade. VMJ strain on agar plates.

On the contrary, the cells of strain SAG 2477 were ellipsoidal to broadly ellipsoidal in shape with broadly rounded ends (Figs 8, 9). Some cells were also slightly bent, but the degree of inclination was not as significant as in the other studied strains. They were 7–16 µm long and 2.5–6.5 µm wide. When cultivated on agar plates, the cell width reached up to 9 µm, and occasional variations of cell shape took on a triangular appearance (Fig. 9). The chloroplast was fragmented, divided into two to four main parts, without a pyrenoid. Older cells contained two or more vacuoles. Reproduction via four and six autospores per autosporangium was observed.

**Fig. 8.**
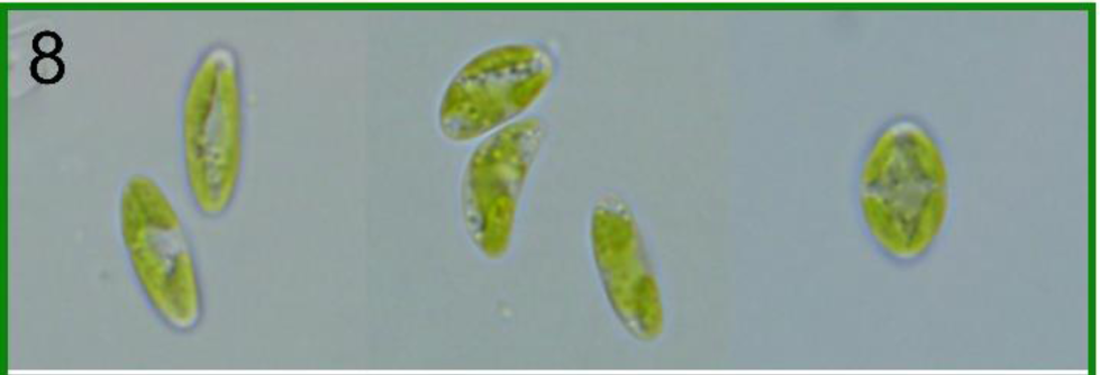
Morphology of the ‘microcroissant’ clade. SAG 2477 in liquid medium.

**Fig. 9.**
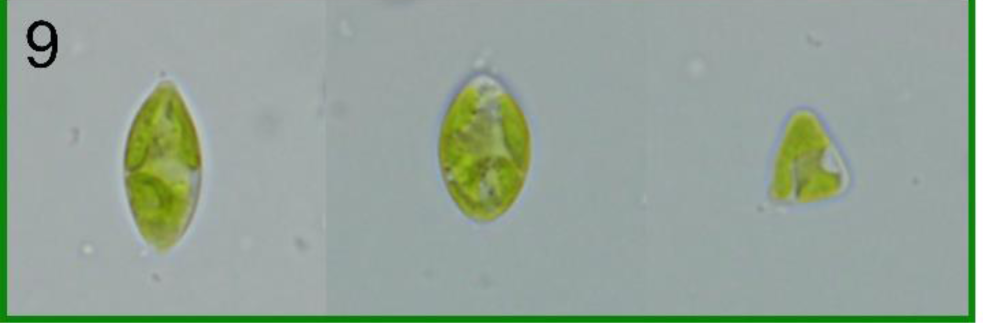
Morphology of the ‘microcroissant’ clade. SAG 2477 on agar plates.

The two additionally studied ACOI strains, 2681 and 3192, both previously identified as *Chlorolobion lunulatum*, demonstrated very similar cell morphologies to our new isolate VMJ (Figs 10–13). More specifically, they both had a crescent-like outline with both ends narrowly rounded or slightly tapered, with the cell length ranging from 7 to 25 µm and the width from 3 to 9 µm. The chloroplast was parietal and trough-shaped and contained a single prominent pyrenoid. Rows of numerous vacuoles were also present within the cells (Figs 11–13; arrows). Reproduction by four or eight autospores per autosporangium was observed. The final carefully inspected strain, *Chlorolobion obtusum* ACOI 732, was found to neatly match the original description of the species by Korshikov (1953) (Figs 14, 15).

**Fig. 10.**
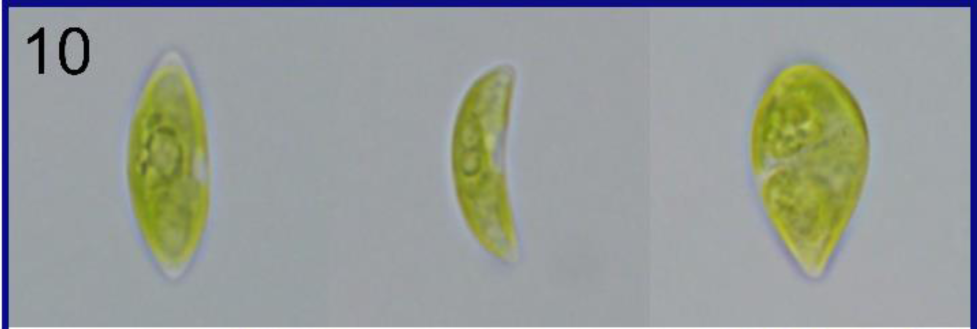
Morphology of the ‘microcroissant’ clade. ACOI 2681 in liquid medium.

**Fig. 11.**
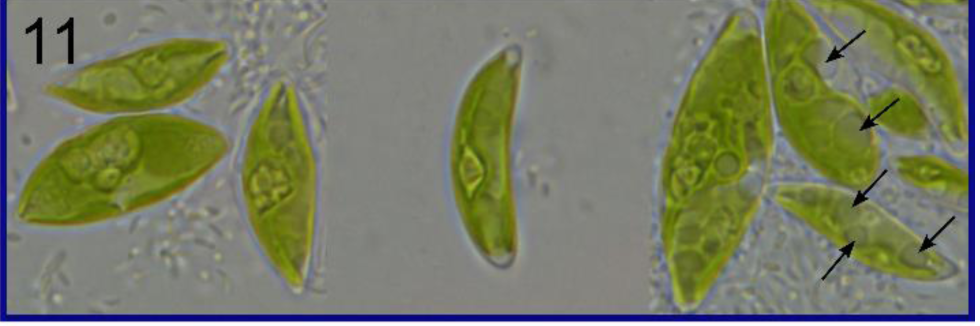
Morphology of the ‘microcroissant’ clade. ACOI 2681 on agar plates.

**Fig. 12.**
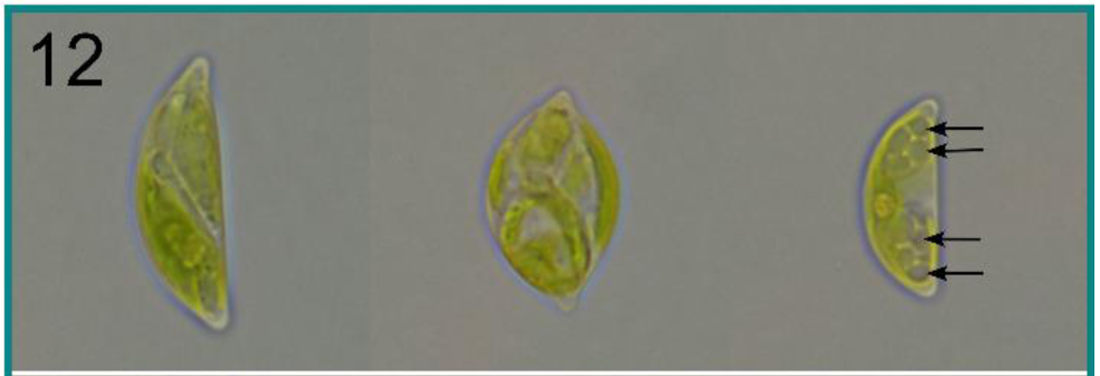
Morphology of the ‘microcroissant’ clade. ACOI 3192 in liquid medium.

**Fig. 13.**
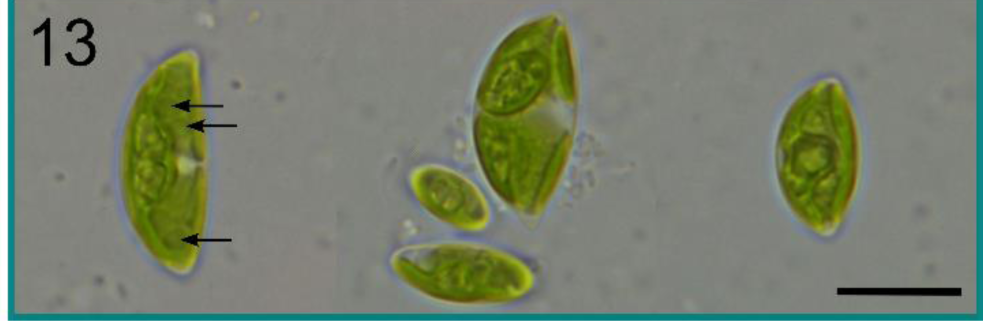
Morphology of the ‘microcroissant’ clade. ACOI 3192 on agar plates. Scale bar = 10 µm.

**Fig. 14.**
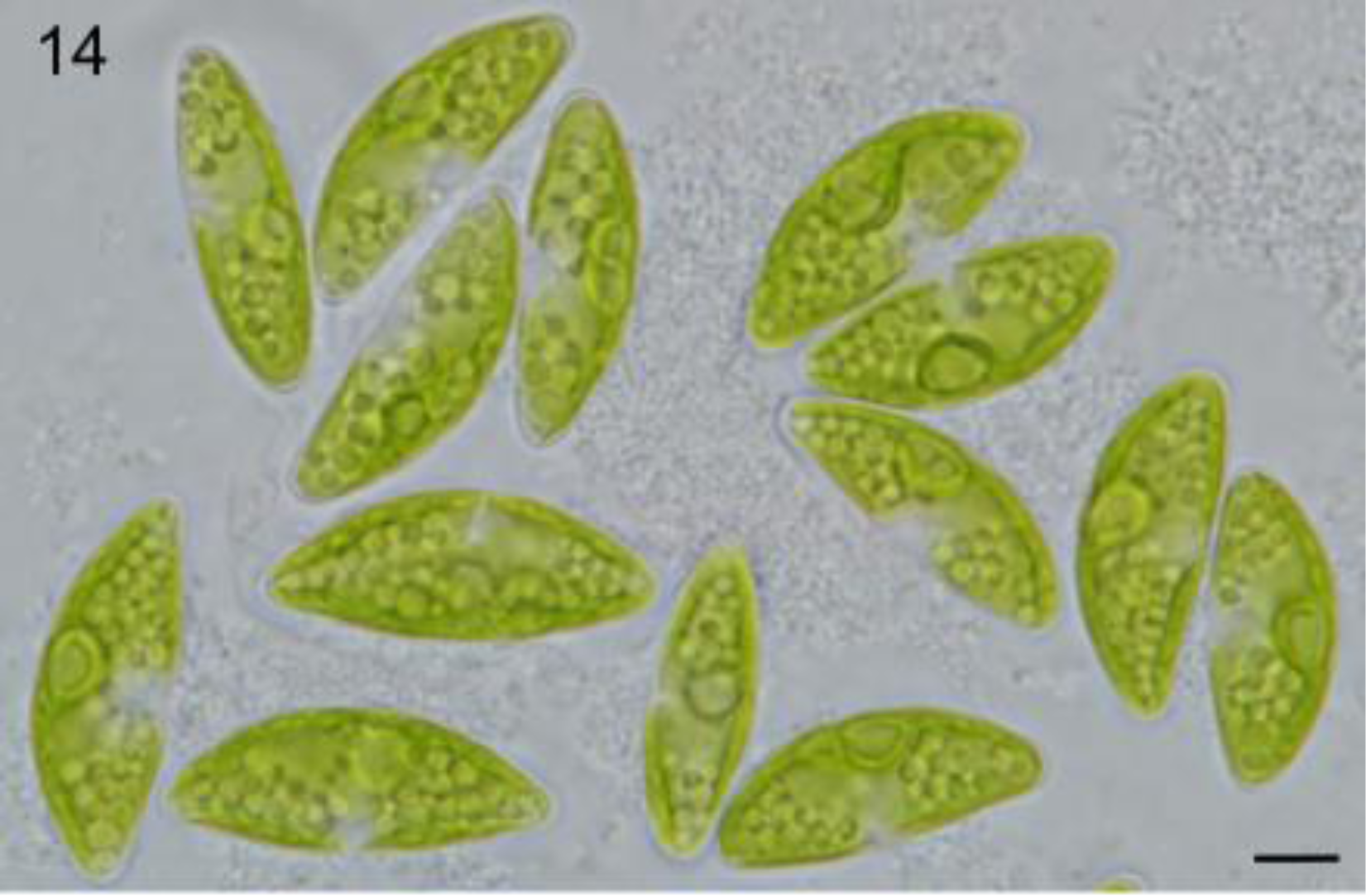
*Chlorolobion obtusum* Korshikov. Original drawings of *Chlorolobion obstusum* from Korshikov (1953).

**Fig. 15.**
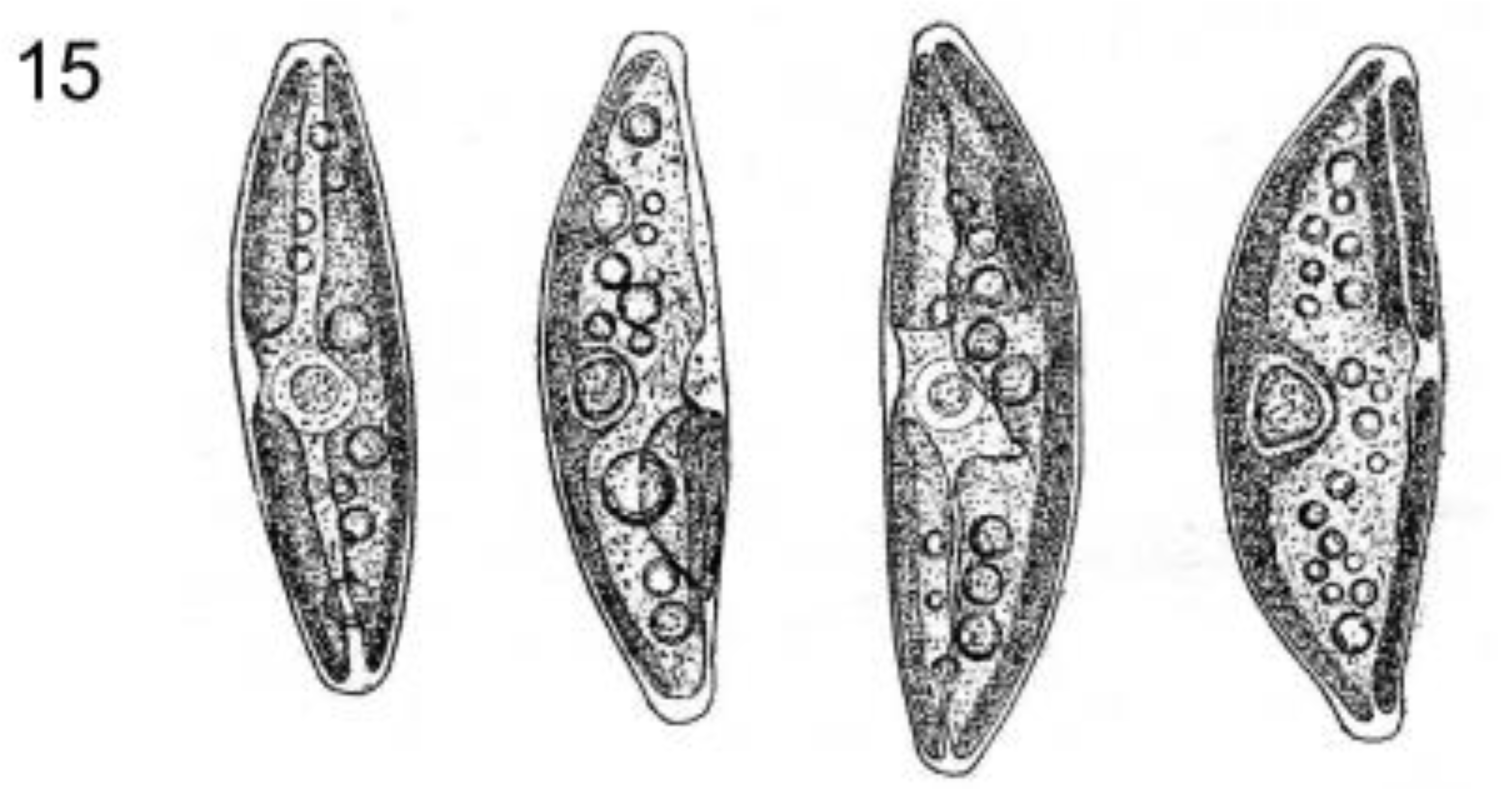
*Chlorolobion obtusum* Korshikov. The matching morphology of ACOI 732. Scale bar = 10 µm.

#### Phylogenetic analyses

To examine the phylogenetic position of the two new isolates (PLY and VMJ), we reconstructed (with ML and Bayesian methods) a 18S rDNA phylogenetic tree including a wide range of Trebouxiophyceae representatives constituting all major lineages of the class. The new isolates were part of a strongly supported clade (ML/BI: 99/1.00) containing several previously studied yet phylogenetically unsettled trebouxiophytes (Fig. 16). Strain PLY was sister to the unidentified desert organism BCP-BC4VF9 sequenced by Fučíková *et al*. (2014) but not studied in further detail, while VMJ branched with the two newly sequenced ACOI strains 2681 and 3192, both isolated from Serra do Gerês in Portugal, and identified as *Chlorolobion lunulatum*. Sequences of ACOI 2681 and 3192 were identical for all three molecular markers sequenced and thus we treat these two strains as conspecific. The VMJ + ACOI 2681/3192 cluster was then sister to the Icelandic *Thorsmoerkia curvula* described by Nicoletti *et al*. (2021) (Fig. 16). All aforementioned organisms constituted a clade further united with strain SAG 2477 as a more deeply diverged sister lineage. This broader clade, which we termed ‘microcroissants’, received high statistical support in both 18S rDNA analyses (ML/BI: 95/1.00). Meanwhile, the last trebouxiophyte strain sequenced in this study, *Chlorolobion obtusum* ACOI 732, did not cluster with ‘microcroissants’ in the 18S rDNA phylogeny but branched instead with *Leptosira* and *Chloropyrula*, with statistical support obtained only in BI analysis (Fig. 16).

**Fig. 16.**
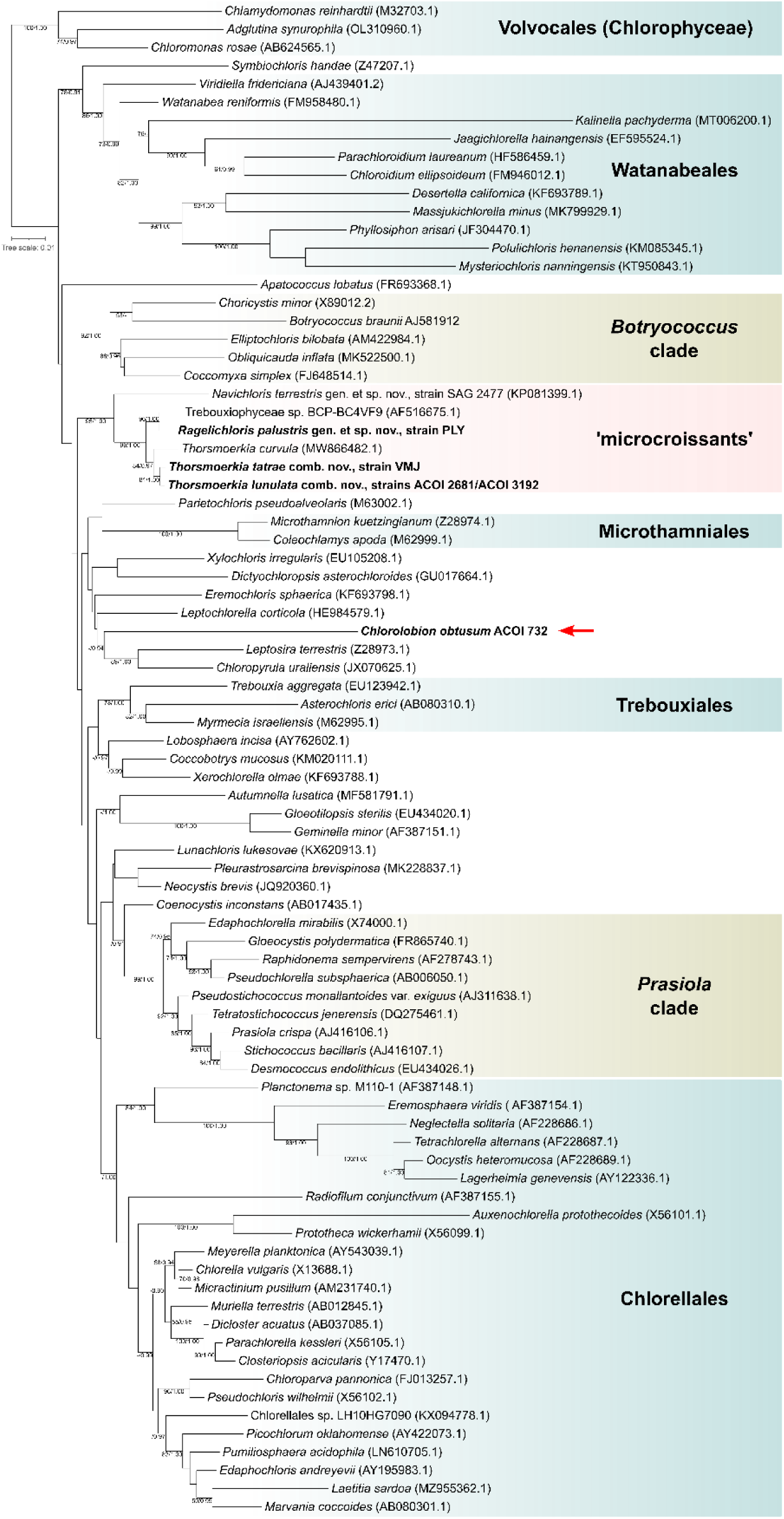
Maximum-likelihood (ML) phylogenetic tree of the class Trebouxiophyceae based on 18S rDNA. Numbers next to branches indicate statistical support values: ML bootstraps/BI posterior probabilities (shown when ML > 50 and BI ≥ 0.90). New sequences are in bold. Red arrow points to the position of the genus *Chlorolobion*. Tree scale indicates substitutions/site.

To further test the grouping of all ‘microcroissants’, we computed a *rbcL*-based phylogenetic tree (Fig. 17). The analysis recovered the same ‘microcroissant’ clade, though only with moderate statistical support in the ML analysis and virtually no support in BI (ML/BI: 70/0.84), consistent with generally lower resolution of the *rbcL* tree compared to the 18S rDNA tree. Crucially, the internal topology of the ‘microcroissant’ clade in the *rbcL* tree was the same as indicated by 18S rDNA sequences. However, in contrast to the 18S rDNA analyses, *Chlorolobion obtusum* ACOI 732 this time branched off as a sister lineage of the ‘microcroissants’ (Fig. 17). Despite the weaker statistical support in ML analysis, BI provided a more convincing evidence for the validity of this relationship (ML/BI: 73/0.99).

**Fig. 17.**
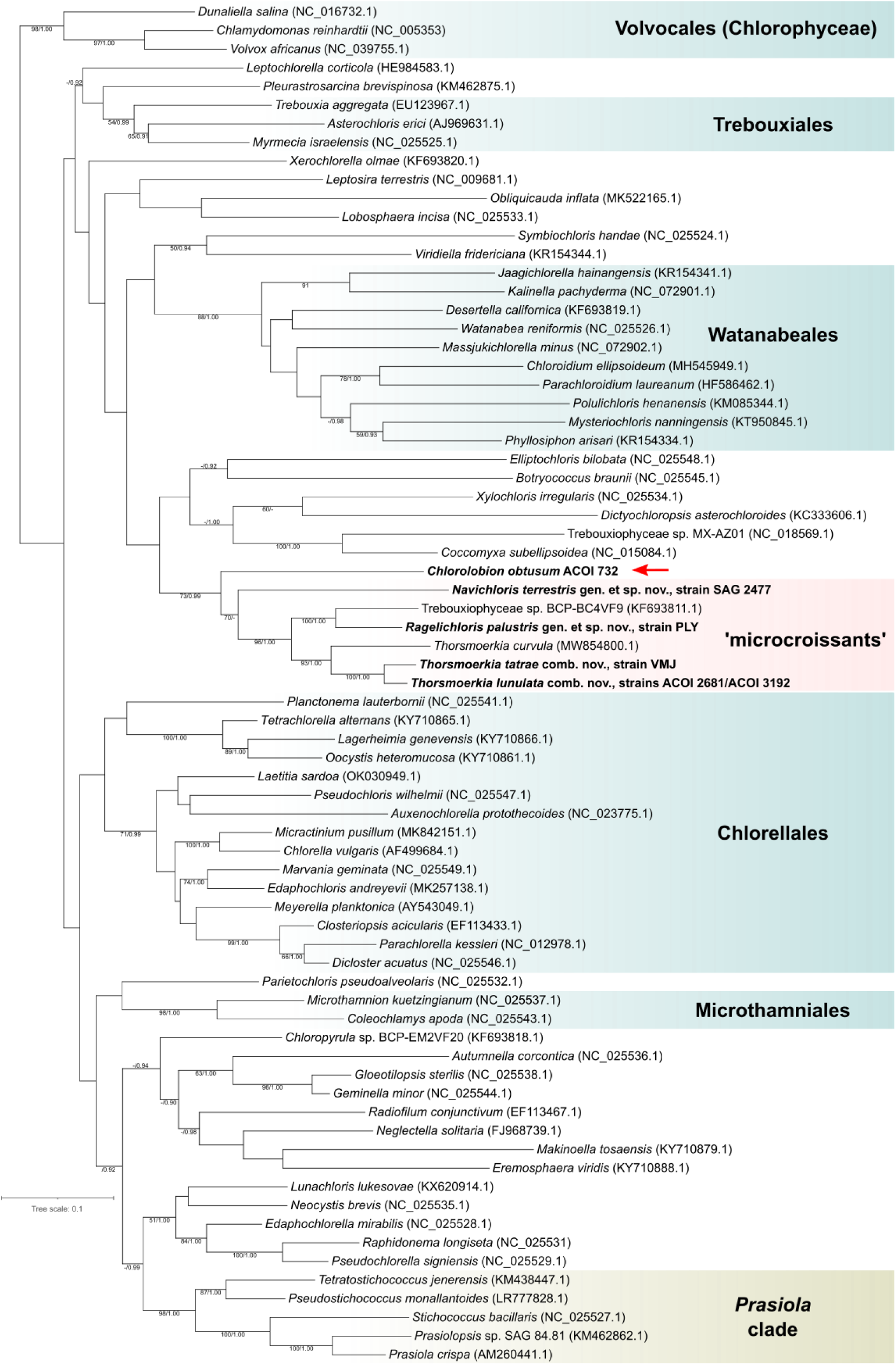
Maximum-likelihood (ML) phylogenetic tree of the class Trebouxiophyceae based on *rbcL* rDNA. The display conventions are the same as for Fig. 16.

Notably, the *rbcL* gene of each VMJ and ACOI 2681/3192 contains a group I intron with an intronic ORF encoding a LAGLIDADG family homing endonuclease. No other case of introns interrupting the *rbcL* gene in any Trebouxiophyceae member was identified by our literature search and inspection of *rbcL* gene sequences available in the GenBank database. (We note that the chloroplast genome sequence ON645925.1, which is assigned in the database to “*Symbiochloris* sp. SG-2018”, i.e. a trebouxiophyte genus, and whose *rbcL* gene is interrupted by two introns, in fact corresponds to the chloroplast genome of the ulvophyte *Symbiochlorum hainanensis*.) The *rbcL* introns of VMJ and ACOI 2681/3192, as well as the proteins encoded by the respective intronic ORFs, are mutually highly similar, consistent with a single acquisition of the intron in the common ancestor of these two microcroissant lineages. Intronic ORFs most similar to those of VMJ and ACOI 2681/3192 come from chloroplast genomes of various non-treboxiophyte green algae and are contained in introns occupying the same position in the *rbcL* gene, but the donor lineage, from which the intron was transferred into the microcroissant subclade, cannot be discerned.

In order to more accurately determine the position of ‘microcroissants’ in the Trebouxiophyceae tree, we used plastome-encoded protein sequences of strain PLY for phylogenomic analysis. The inferred phylogenetic tree placed PLY as a sister lineage to the coccoid subaerial alga *Xylochloris irregularis*, with the two having the filamentous soil alga *Leptosira terrestris* as their immediate relative. The close phylogenetic relationship among these three algae received full statistical support. Moreover, the clade comprising *Leptosira*, *Xylochloris*, and strain PLY emerged with robust support (ML: 93%) as a group sister to Microthamniales (Fig. 18).

**Fig. 18.**
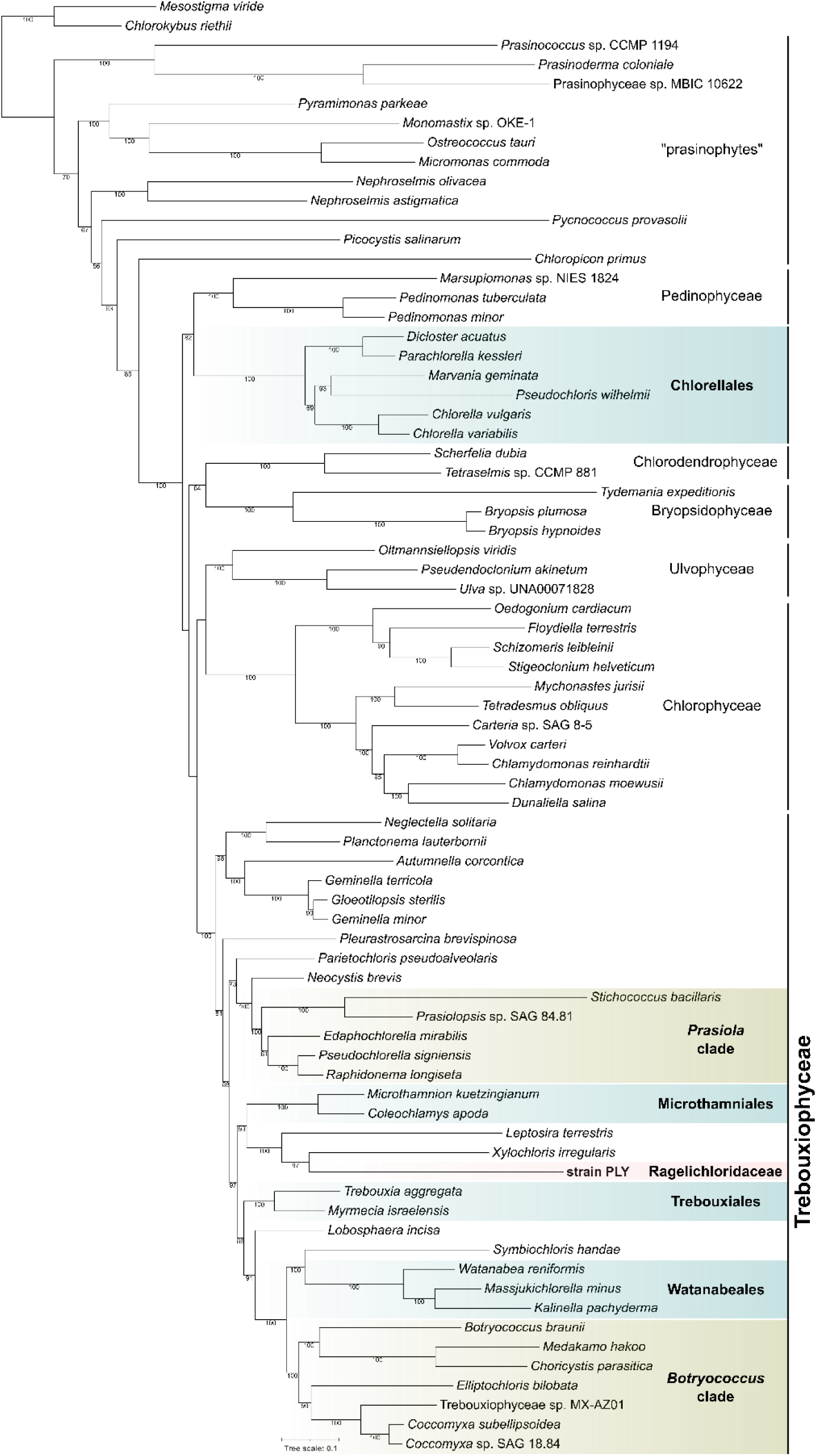
Maximum-likelihood (ML) phylogenetic tree inferred from a concatenated data set of 79 plastome-encoded proteins from the green algae. The alignment used for the tree inference consisted of 16,700 amino acid positions. Bootstrap support values shown when > 50. Tree scale indicates substitutions/site.

### ITS2 rDNA secondary structures

To gain additional insights into the genetic diversity and relationships of ‘microcroissants’, we sequenced the ITS2 regions of the respective algae and predicted their secondary structures. Secondary structure analysis allows comparison of the base-pair interactions and their compensatory mutations (=compensatory base changes; CBCs) or base substitutions on only one side of the stem structures (hemi-CBCs) between close relatives. As illustrated in Fig. 19, the most striking difference among the examined stains was the absence of a typical helix IV motif in VMJ, ACOI 2681/3192, and *Thorsmoerkia curvula* (MW866482.1), i.e. all representatives of a particular subclade of ‘microcroissants’ phylogenetic analyses (Figs 16, 17). In addition, ACOI 2681/3192 lacked a short side loop in helix III, a feature that was present in the two closest relatives, making the ITS2 secondary structure of this strain pair the most distinctive one within the ‘microcroissant’ clade. A detailed comparison of VMJ and ACOI 2681/3192 revealed the presence of a single CBC in helix I and two hemi-CBCs in helix III between the two strains, while comparison with *Thorsmoerkia curvula* (MW866482.1) displayed a total of three CBCs distributed across the helices I and II as well as in the 5.8S and 28S stem in both strains. The comparison of two *Sphagnum*-associated strains, PLY and VMJ, showed five CBCs (one in the helix I, three in the helix III, and one in the stem) and four hemi-CBCs. Interestingly, the ITS2 secondary structure of strain PLY exhibited a closer resemblance to that of the more distantly related SAG 2477 compared to the closer relatives (Fig. 19; the ITS2 sequence from BCP-BC4VF9, the closest known relative of PLY, is not available). They exhibited six CBCs and three hemi-CBCs between themselves. The highest number of CBCs (eight) were found when comparing strain PLY with ACOI 2681/3192 and *Thorsmoerkia curvula*. As expected, the ITS2 secondary structure of ACOI 732 was the most different one (Fig. 19). An UPGMA cladogram, based on genetic distances within the sequence + structure alignment of the ‘microcroissant’ clade (with ACOI 732 as an outgroup), exhibited topology agreeing with strain similarities evident from the graphical visualisation of the ITS2 structural models (Fig. 19).

**Fig. 19.**
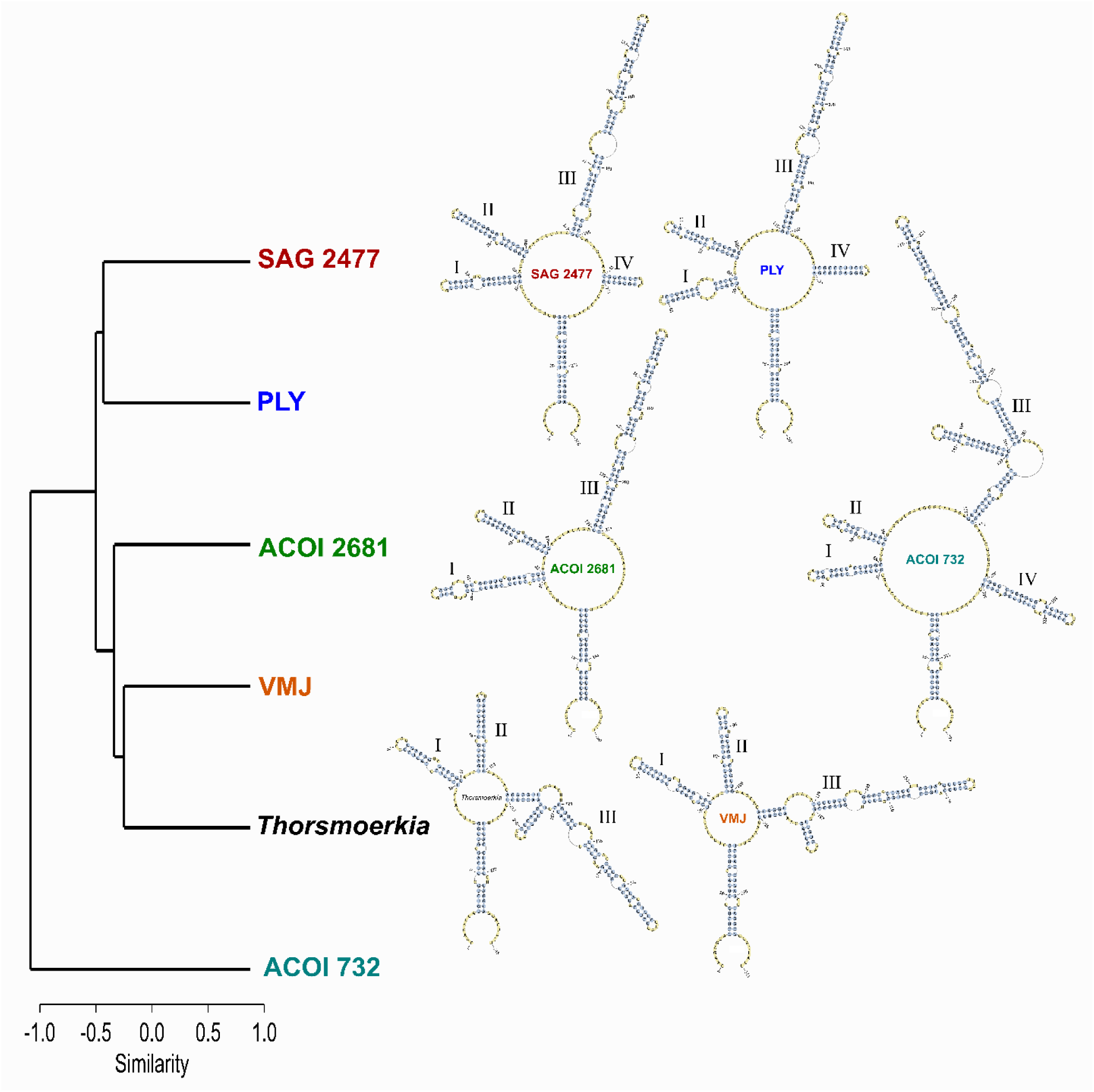
The UPGMA cladogram based on genetic distances within the sequence + structure alignment of the ‘microcroissant’ clade and ITS2 secondary structures of its members. Nucleotides that are paired are shown in blue, while nucleotides that are unpaired are shown in yellow. Helices are marked I to IV.

## Discussion

### Identification of the new strains

Morphologically the two newly isolated strains (PLY and VMJ) immediately resemble members of the family Selenastraceae (Chlorophyceae) (Krienitz *et al*., 2001). Examples of particularly similar selenastracean algae are different species of the genus *Monoraphidium* Komárková-Legnerová, including *M. terrestre*, *M. dybowskii*, and *M. tatrae*. However, some minute differences do exist. For example, *M. dybowskii* and *M. terrestre* have a pyrenoid, with and without a starch envelope, respectively, and the latter species is also characterized by a prominent secretion of mucilage (Krienitz *et al*., 2001). Strain VMJ also contains a starch-covered pyrenoid as evidenced by the light microscopy (Fig. 7), but it does not produce cylindrical cells as *M. dybowskii*, and the maximum cell length measured for VMJ considerably exceeds that of *M. dybowskii*. In contrast, strain PLY did not show the presence of the pyrenoid matrix even under TEM (Figs 2, 3, 5), and we did not detect any mucilage secretion in any of the two newly isolated strains. Most crucially, the position of the two discussed *Monoraphidium* species in Selenastraceae has been confirmed by molecular means (Krienitz *et al*., 2001).

Among the species mentioned, strain VMJ most closely resembles *Monoraphidium tatrae* in terms of size, cell and chloroplast shape, the presence of the two prominent vacuoles in some cells, as well as the presence of multiple smaller vacuoles in others. Thanks to the available detailed line drawings, we could immediately confirm all morphological matches of our strain to *M. tatrae* (Komárek & Fott, 1983). The only significant discrepancy between *M. tatrae* and strain VMJ is the absence of the pyrenoid in the former. We could link our isolate VMJ to *M. tatrae* only because the presence of the pyrenoid was not obvious in all cells when the strain was cultivated in liquid medium; Fig. 6). Noteworthy, *M. tatrae* was originally described as *Chlorolobion tatrae* (Hindák, 1970) and only later on was moved to *Monoraphidium* on the grounds of the absence of the pyrenoid (Hindák, 1977). It is therefore significant to note that many of the *Monoraphidium* species described as pyrenoid-less in the past have been shown to actually contain a pyrenoid after a thorough culture-based re-investigation (Krienitz *et al*., 2001). Such discrepancies clearly suggest that the presence or absence of the pyrenoid in the original descriptions of green algae should be taken with caution, especially for the minute ones. *Chlorolobion tatrae* was described from a field sample, so it is not at all surprising that the pyrenoid could have been overlooked. In our case, the pyrenoid was consistently observed in strain VMJ only when grown on agar plates (Fig. 7). Considering strain VMJ as *C. tatrae* is also hinted to by the shared habitat of shallow waters in high elevations. Specifically, *C. tatrae* was found in shallow pools in the area of mountain lakes (Päť Spišských plies) in High Tatras (Slovakia) (Hindák, 1970), while VMJ was isolated from a *Sphagnum*-dominated shore of a lake (Velké mechové jezírko) in Jeseníky Mountains, Czech Republic.

Unlike strain VMJ, we were unable to confidently match our second isolate, PLY, to any previously reported species. Apart from *Monoraphidium*, other carefully checked genera included *Podohedra* Düringer and *Keratococcus* Pascher (Komárek & Fott, 1983). However, *Podohedra* has a long stalk, and *Keratococcus* typically contains long spiny projections at both ends, features that are absent in strain PLY. We do accept the possibility that the organism could have been found and described before, but linking it to any of the existing name is extremely challenging considering the widespread uniformity (the lack of species-specific features) of the croissant-like morphotype, further complicated by phenotypic plasticity (Figs 1, 4). In order to avoid any confusion and misidentification, we find it most reasonable to establish strain PLY as a novel taxon with a clearly defined phylogenetic position and reference sequences (see below for Formal taxonomy).

### *Chlorolobion* Korshikov belongs to Trebouxiophyceae

Our identification of strain VMJ as *C. tatrae* prompted further investigation into the genus *Chlorolobion* and where it falls in the green algal phylogeny. The genus was first described by Korshikov (1953) with the type species *Chlorolobion obtusum*. Later, Hindák (1970) provided formal descriptions of additional two species, *C. tatrae* and *C. lunulatum*. In the course of time, five other species (*C. braunii* (Nägeli) Komárek, *C. glareosum* (Hindák) Komárek, *C. saxatile* (Komárková-Legnerová) Komárek, *C. guanense* Comas, and *C. tianjinensis* Wang & Feng) were assigned to the genus (Komárek & Fott, 1983; Guiry & Guiry, 2023). Among the species described, only *Chlorolobion braunii* has been subjected to a molecular taxonomic investigation and phylogenetic analysis revealed its placement within Selenastraceae (Garcia da Silva *et al*., 2017). However, the needle-like morphology of *C. braunii* and some other nominal *Chlorolobion* species differs quite substantially from the morphology of the type species *C. obtusum* (Komárek & Fott, 1983), raising the possibility *Chlorolobion* as currently circumscribed as a polyphyletic unit.

Thus, finding the general morphological similarity of our isolates to the “typical” *Chlorolobion* species, we found it crucial to study additional strains identified as members of *Chlorolobion* and available from the ACOI culture collection (Table 1) in an attempt to clarify their possible relationship to the ‘microcroissant’ clade. Of the nine such strains investigated, six were uncovered to represent members of the class Chlorophyceae, specifically the family Selenastraceae, based on complete or partial 18S rDNA sequences (data not shown), but three of them, ACOI 732, 2681, and 3192, turned out to be trebouxiophytes, representing two independent lineages (Figs 16, 17). Of crucial significance was the strain ACOI 732 isolated and identified by the ACOI curator M. F. Santos as representative of *Chlorolobion obtusum* – the type species of the genus. Upon examining the strain’s morphology, we endorse the validity of this identification (Figs 14, 15). By demonstrating the position of *C. obtusum* as an independent lineage in Trebouxiophyceae we validate the separate generic status of *Chlorolobion* and clarify its assignment into a particular green algal class. The generic assignment of other nominal *Chlorolobion* species requires further scrutiny. Considering our results, some species may need to be reassigned to a different genus or genera in Selenastraceae, but a separate careful reassessment of the respective strains, ideally as part of a broader critical revision of the whole family Selenastraceae, is needed. Our results, however, do provide a strong case for a taxonomic revision of *C. tatrae* and *C. lunulatum*, which is presented in the next section.

### Expanding the genus *Thorsmoerkia* and introducing *Ragelichloris* gen. nov

As revealed by our molecular phylogenetic analyses, the ACOI strains 2681 and 3192 are closely related to our VMJ isolate. It is thus of high significance that both ACOI strains were identified as *Chlorolobion lunulatum*, solidifying our independent identification of VMJ strain as a closely related species, *C. tatrae*. Indeed, all the morphological features exhibited by ACOI 2681 and 3192 strains are consistent with those of *C. lunulatum* (Figs 10–13). The other ACOI strains assigned by the culture collection to *C. lunulatum* (231, 811) represent selenastracean algae (Table 1) and were obviously misidentified, because their morphology does not match the description of the species (Fig. S3). Even though Komárek and Fott (1893) suggested possible conspecificity of *C. tatrae* and *C. lunulatum*, we do not endorse it. Both VMJ and the two ACOI strains are morphologically very similar, but phylogenetic divergence (especially evident in the *rbcL* phylogeny) and significant differences in their ITS2 rDNA secondary structures (Fig. 19) support their classification as distinct species. Needless to say, *C. tatrae* and *C. lunulatum* cannot stay in the genus *Chlorolobion* as they are not specifically related to *C. obtusum*.

All molecular markers employed in this study instead consistently place *Chlorolobion tatrae* and *C. lunulatum* as relatives of *Thorsmoerkia curvula*, the type and presently the only species of the recently established genus *Thorsmoerkia* Remias & Procházková (Figs 16, 17). It thus seems reasonable to resolve the untenable generic classification of the two nominal *Chlorolobion* species by expanding the circumscription of the genus *Thorsmoerkia* to embrace these two species as new combinations *Thorsmoerkia tatrae* and *Thorsmoerkia lunulata* (see Formal taxonomy).

It is then a matter of an arbitrary decision whether strain PLY should also be a part of *Thorsmoerkia* or not. The former treatment would be compatible with the general morphological similarity of strain PLY and *Thorsmoerkia* spp., but the phylogenetic distance between strain PLY and *Thorsmoerkia* is comparable to that observed between different genera within the Chlorellales or the *Prasiola*-clade (Figs 16, 17), making a case for classification of strain PLY as a separate genus. In addition, compared to the three *Thorsmoerkia* species, whose ITS2 secondary structure lacks helix IV, strain PLY has retained the plesiomorphic state with helix IV, making its ITS2 structure more similar to that of SAG 2477, which is even more distantly related to *Thorsmoerkia* than strain PLY (Fig. 19). In addition, strain PLY exhibits more tapered cell ends and lacks rows of vacuoles, the latter feature so characteristic for *Thorsmoerkia* species (Figs 1, 2, 6, 7, 10–13). We thus find it reasonable to place strain PLY into a genus separate from *Thorsmoerkia*, and as there seems to be no other previously described genus that could be considered as a taxonomic home of strain PLY, we describe it as a new species in a new genus, *Ragelichloris palustris* (see Formal taxonomy). Based on our phylogenetic analyses, the now-lost strain BCP-BC4VF9 found in a Mexican desert (Fučíková *et al*., 2014) is related to *R. palustris* and could be naturally classified in the same genus, yet clearly as a different species.

### *Navichloris terrestris*: a new genus and species of Trebouxiophyceae

Touching on strain SAG 2477, there are similarities with other terrestrial coccoid algae, including certain species the genus *Coccomyxa* Schmidle and *Chlorocloster* Pascher. The former is confidently a trebouxiophyte, yet unrelated to SAG 2477, whereas the latter was described as a member of Pascher’s “Heterokonten” and without evidence *contra* it is by default presently classified in a descendant of the traditionally circumscribed heterokont algae, i.e. in the ochrophyte class Xanthophyceae. However, numerous former members of this grouping have proved to belong to different major lineages of eukaryotes, including green algae (e.g. Eliáš *et al*., 2013). Indeed, one of the nominal *Chlorocloster* species, *Chlorocloster engadinensis* Vischer, has been formally transferred to the trebouxiophyte genus *Chloroidium* upon scrutiny with modern methods (Darienko *et al*., 2010). We were, therefore, compelled to critically compare SAG 2477 with *Chlorocloster*, typified by *Chlorocloster terrestris*, which was described as a common terrestrial alga occurring in meadow and forest soil (Pascher, 1925). Indeed, both organisms share the same habitat, cell size, and the illusion of the presence of three chloroplasts within the cell. However, a significant difference between them is that the cells of *C. terrestris* have clearly narrowed ends and also exhibit an S-shape, while the cell termini of SAG 2477 are rounded and the alga is not noticeably S-shaped. Another similar species is *Chlorocloster simplex* Pascher also matching the general appearance of the ‘microcroissant’ but typically having very unequal ends (Pascher, 1938). Finally, the ellipsoidal to oval cells and highly fragmented chloroplast of strain SAG 2477 also bear a resemblance to another putative xanthophyte genus *Ellipsoidion* Pascher, including the species *E. simplex*, *E. stichococcoides*, *E. acuminatum*, and *E. pulchum*. Because of the morphological variability exhibited by strain SAG 2477 (Figs 8, 9), it is challenging to assign it to a specific *Ellipsoidion* species. It is though worth noting that one of the former *Ellipsoidion* species, *E. parvum*, has been reclassified as a conspecific member of the trebouxiophyte alga *Neocystis brevis* (Eliáš *et al*., 2013). Therefore, a close relationship between strain SAG 2477 and certain *Ellipsoidion* species would not be unexpected. However, as for now, considering the deeply branching nature of the SAG 2477 from the other studied strains (*Ragelichloris* and *Thorsmoerkia*) along with its unmatched morphology with the previously described species, we place the SAG 2477 into a new genus and species, *Navichloris terrestris*, expanding thus the taxonomic diversity of trebouxiophyte algae (see Formal taxonomy).

### ‘Microcroissants’ constitute a higher-level clade of Trebouxiophyceae

By employing the conservative molecular markers 18S rDNA and *rbcL* we obtained consistent support for the monophyly of a group encompassing the three genera, *Navichloris*, *Ragelichloris*, and *Thorsmoerkia* (Figs 16, 17). The phylogenetic divergence of the ‘microcroissants’ (i.e. *Navichloris* + *Ragelichloris* + *Thorsmoerkia*) clade from other trebouxiophytes is comparable even to that of different formally delineated orders within the class. In addition, the similar croissant-like morphology of the three genera also suggest their assignment to the same higher-level taxonomic rank. As for now, we interpret the ‘microcroissants’ clade as a new family-level taxon, which we describe as Ragelichloridaceae (see Formal taxonomy).

Chloroplast phylogenomics, which surpasses the better-sampled single-gene phylogenies in terms of resolution of deeper branches of the trebouxiophyte phylogeny, uncovered Ragelichloridaceae, represented by *Ragelichloris palustris* (strain PLY), as part of a broader clade additionally including the genera *Xylochloris* and *Leptosira* (Fig. 18). The latter two genera were found to be related to each other in a previous less-well sampled chloroplast phylogenomic tree and together denoted as the “*Xylochloris* clade” (Lemieux *et al*., 2014). Our results thus expand this emerging grouping by adding Ragelichloridaceae. Considering the results of the 18S rDNA phylogeny (Fig. 16), the genera *Dictyochloropsis* (potentially related to *Xylochloris*; Figs 16, 17) and *Chloropyrula* (specifically related to *Leptosira*; see also Gaysina *et al*., 2013) are also candidate members of the *Xylochloris* clade, which likely holds also for *Chlorolobion* based on the *rbcL* tree (Fig. 17). The *Xylochloris* clade thus seems to represent a candidate new order in Trebouxiophyceae.

We refrain from a formal definition of the new order in this study and advocate for further phylogenomic and phylotranscriptomic studies to test the aforementioned inferences on the taxonomic composition of the putative order. Nevertheless, we point to a potential synapomorphy of this grouping, namely the absence of the *ycf12* gene previously noticed to be shared by the chloroplast genomes of *Xylochloris* and *Leptosira* (Turmel *et al*., 2015). The gene encodes a component (also known as Psb30) of the photosystem II complex (Inoue-Kashino *et al*., 2011) and is virtually omnipresent in chloroplasts of green algae and plants, with additional exceptions found only among seed plants (all angiosperms and a subset of gymnosperms) (Turmel & Lemieux, 2018; Kwon *et al*., 2020). In most trebouxiophytes, including taxa in the phylogenetic vicinity of the *Xylochloris* clade (Microthamniales, Trebouxiales), the gene is part of a gene block *psbK-ycf12-psaM*. Our *R. palustris* transcriptome assembly contains a transcript that includes the *psaM* gene directly downstream of *psbK*, with no *ycf12* in between them (Fig. S4), paralleling thus the situation in the fully sequenced chloroplast genomes of *Xylochloris irregularis* (NC_025534.1) and *Letosira terrestris* (NC_009681.1). It is thus likely that *R. palustris* shares the *ycf12* loss with the two previously investigated members of the *Xylochloris* clade. Indeed, no *ycf12* homolog could be found in the whole transcriptome assembly *R. palustris*, and it is missing also from the transcriptome assembly available for *Leptosira terrestris*. This indicates that the *ycf12* loss from the chloroplast genome in the *Xylochloris* clade has not been compensated for by the transfer of the gene to the nuclear genome, and that the photosystem II of these algae differs from that of other green algae by the absence of the *ycf12*-encoded subunit.

### Formal taxonomy

#### Ragelichloridaceae Barcytė, fam. nov

DESCRIPTION: Unicellular croissant-shaped or broadly ellipsoidal cells with rounded ends. Solitary or in loose groups. Uninucleate. Chloroplasts are one per cell and lobed in older cells, with or without a pyrenoid. With or without vacuoles. Asexual reproduction by autospores. Found in freshwater and terrestrial habitats.

TYPE GENUS: *Ragelichloris* Barcytė

REMARKS: The family as delimited here presently includes *Ragelichloris* gen. nov., *Navichloris* gen. nov., and *Thorsmoerkia* Remias & Procházková.

#### *Ragelichloris* Barcytė, gen. nov

DESCRIPTION: Unicellular, solitary or in loose groups, morphologically plastic algae demonstrating croissant-like to drop-like shapes with narrowly-rounded ends. Uninucleate. Chloroplast single, parietal, trough-shaped, or multilobed. Asexual reproduction by autospores. The genus differs from other genera in nuclear 18S and ITS2 rDNA and chloroplast *rbcL* sequences.

#### TYPE SPECIES: *Ragelichloris palustris* Barcytė, sp. nov

ETYMOLOGY: The genus name *Ragelichloris* is derived from the Lithuanian word "ragelis" meaning croissant and Greek word *khloros/χλωρός* meaning "pale green".

#### *Ragelichloris palustris* Barcytė, sp. nov. (Figs 1–5)

DESCRIPTION: In liquid cultures cells are croissant-shaped, 12–20 µm long and 3–8 µm wide; on agar plates cells are drop-like. Chloroplast parietal and trough-shaped, divided in several parts in older cells, without a pyrenoid. Reproduction by 2-4-8 autospores, liberated by rupture of sporangial cell wall. Sexual reproduction not observed.

HOLOTYPE: TEM block (cells in resin) deposited at the CAUP culture collection (Charles University, Prague) with ID XXXX.

TYPE LOCALITY: Raised bog Plynoja, Pagramantis Regional Park, Tauragė, Lithuania (55°19′23″ N, 22°8′18″ E).

ETYMOLOGY: The Latin word "*palustris*" means "of the marsh or swamp" or "marshy, swampy". It points to the fact that the species was isolated from the peat bog.

#### *Navichloris* Hodač & Barcytė, gen. nov

DESCRIPTION: Vegetative cells solitary, ellipsoidal to broadly ellipsoidal. Uninucleate. The single chloroplast is parietal, fragmented. Asexual reproduction via autospores. The genus differs from other genera in nuclear 18S and ITS2 rDNA and chloroplast *rbcL* sequences.

TYPE SPECIES: *Navichloris terrestris* Hodač & Barcytė

ETYMOLOGY: Genus name comes from a Latin word *navis* meaning "ship", emphasizing ship-shaped cells and Greek word *khloros*/*χλωρός* meaning "pale green"’.

#### *Navichloris terrestris* Hodač & Barcytė, sp. nov. (Figs 8, 9)

DESCRIPTION: In liquid cultures cells are solitary, ellipsoidal to broadly ellipsoidal with broadly rounded ends, 7–16 µm long and 2.5–6.5 µm wide; on agar plates irregular shapes may appear. Can be slightly bent. Chloroplast single, parietal, and divided into two to four main parts. Pyrenoid absent. Vacuoles may be present. Asexual reproduction via 4-6 autospores. Sexual reproduction not observed.

HOLOTYPE: Metabolically inert (cryopreserved) culture SAG 2477 at the Culture Collection of Algae at the University of Göttingen, Germany (SAG).

TYPE LOCALITY: Soil in spruce forest, Swabian Alb, Germany (48° 24’ 44.145" N, 9° 21’ 20.127" E).

ETYMOLOGY: The species name "*terrestris*" comes from the Latin word "*terra*" meaning "earth" or "soil", emphasizing that this species is isolated from soil.

#### *Thorsmoerkia tatrae* (Hindák) Barcytė, comb. nov. (Figs 6, 7)

BASIONYM: *Chlorolobion tatrae* Hindák, 1970, Algol. Stud. 1: 7–32.

SYNONYMS: *Monoraphidium tatrae* (Hindák) Hindák, 1977: 109.

#### *Thorsmoerkia lunulata* (Hindák) Barcytė, comb. nov. (Figs 10, 11)

BASIONYM: *Chlorolobion lunulatum* Hindák, 1970, Algol. Stud. 1: 7–32.

## Acknowledgements

This article has been produced with the financial support of the European Union under the LERCO project number CZ.10.03.01/00/22_003/0000003 via the Operational Programme Just Transition and of the Czech Science Foundation project 23-06203S.

## Supplementary information

**Table S1.** Thermal profiles of PCR reactions.

**Table S2.** GenBank accession numbers of chloroplast genomes used for phylogenomic analysis.

**Fig. S1.** Bog Plynoja where strain PLY was isolated from.

**Fig. S2.** The Great Moss Lake (Velké mechové jezírko) where strain VMJ was isolated from.

**Fig. S3.** Light micrographs of the studied ACOI strains that do not belong to the Trebouxiophyceae class.

**Fig. S4**. *Ragelichloris palustris* PLY polycistronic transcript containing coding sequences of the *psbK* and *psaM* genes, but lacking the *ycf12* coding sequence in between them.

## Author contributions

D. Barcytė: original concept, microscopy, phylogenetic analyses, drafting and editing manuscript; L. Hodač: analysis of molecular data, drafting and editing manuscript; M. Eliáš: funding acquisition, drafting and editing manuscript.

