## Supplementary figures S1-S4 for "Freshwater ‘microcroissants’ shed light on a novel higher-level clade within Trebouxiophyceae and reveal the genus *Chlorolobion* as a trebouxiophyte"

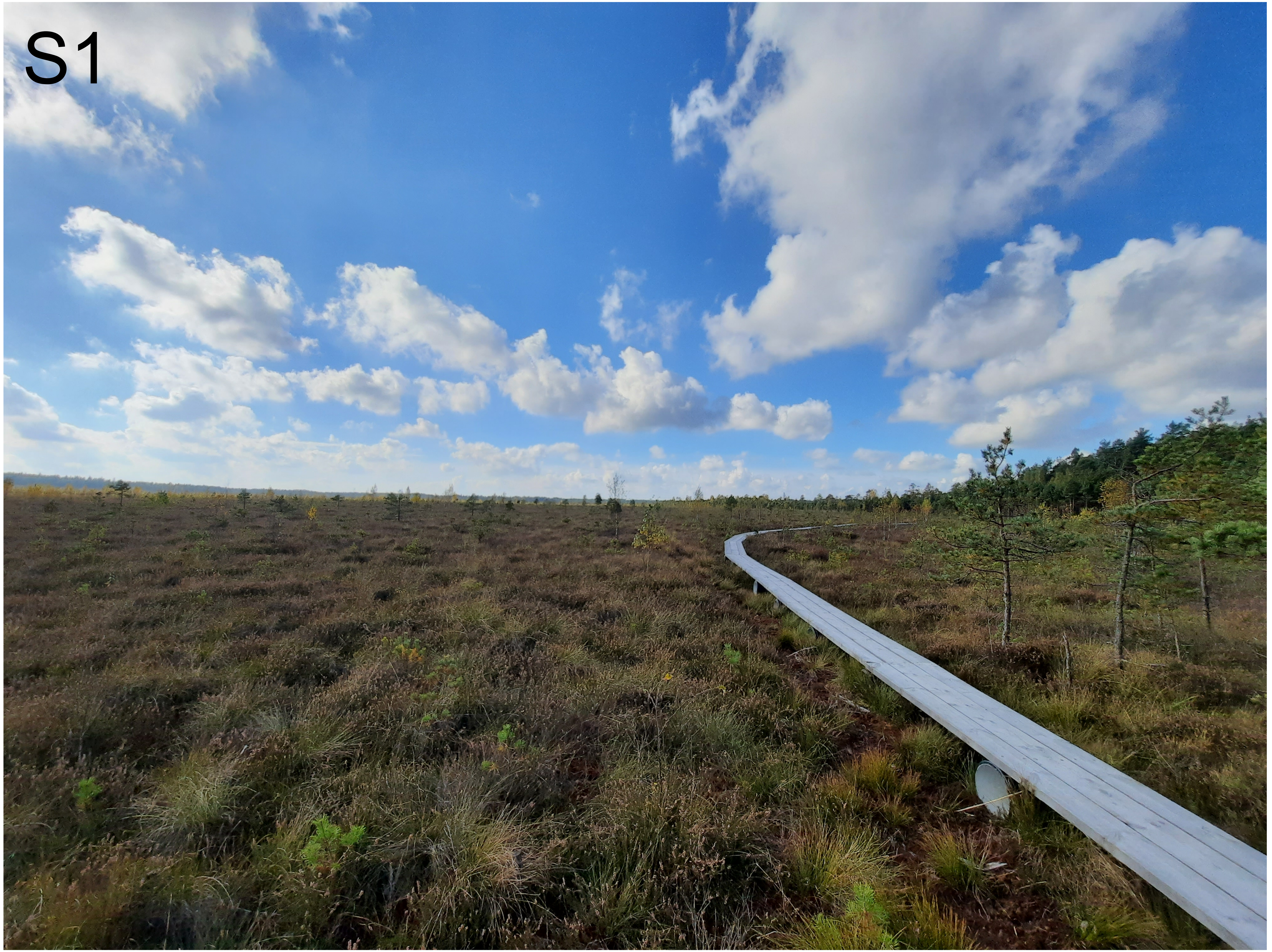


Fig. S1. Bog Plynoja where strain PLY was isolated from.

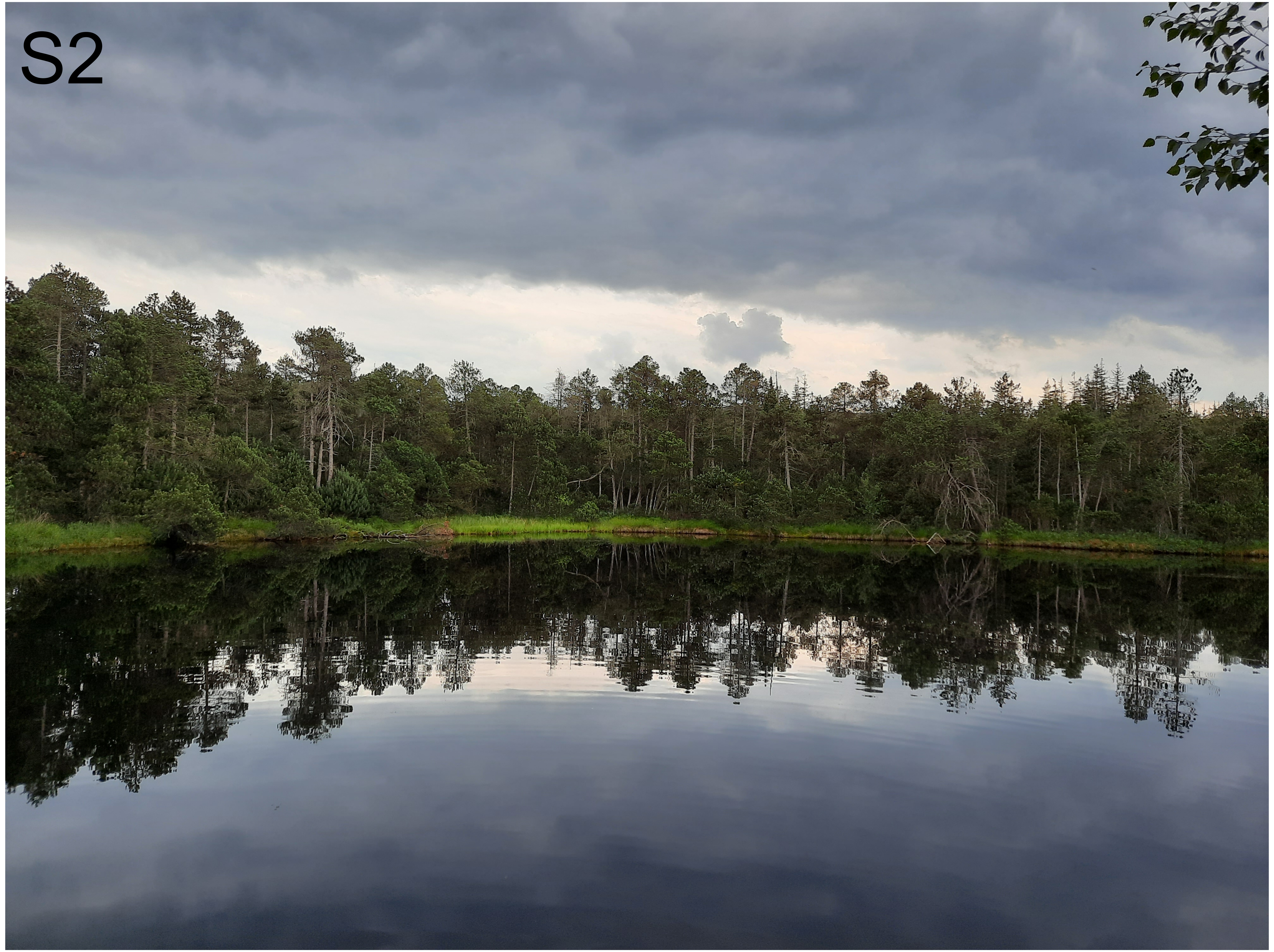


Fig. S2. The Great Moss Lake (Velké mechové jezírko) where strain VMJ was isolated from.

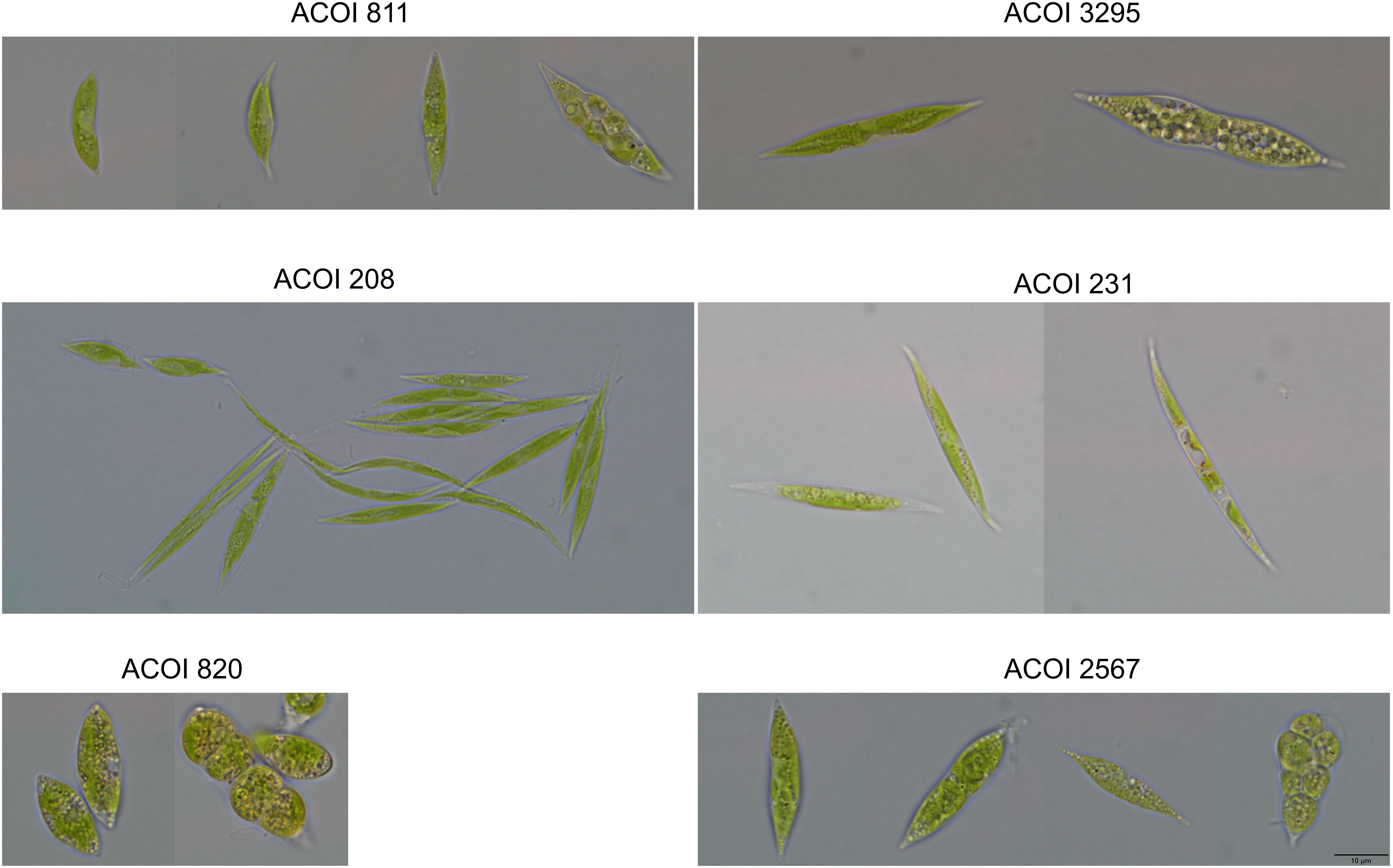


Fig. S3. Light micrographs of the studied ACOI strains that do not belong to the Trebouxiophyceae class.

**Fig. S4.** *Ragelichloris palustris* PLY polycistronic transcript containing coding sequences of the *psbK* and *psaM* genes, but lacking the *ycf12* coding sequence in between them.

```
>TRINITY_DN36743_c0_g1_i1 len=2314 path=[2292:0-2313] [-1, 2292, -2]
AGCCGCCCTAAGGTCGTAAGGCCTGACACAACGACCTTGTGTAAGAGGGGGCTCGAAGGC
TAACAAATCTTTGATTTGTAACCCCGGCCATCGAGGGGGTTTGTAAAATTCATAAAAGAG
GTTACCTTATTGCCTTAGTAAATAAAAGTAGAGTTTTTTAGAAAGATAAAAGTTCTATCA
ATTCTTCAAAGTTTTTTAATGATAATCAAAAATGTGTAATATATCTTTATAGCCTTTACC
TTAATTTTTGAGTTATCTTTTCGTATTAAGAGGCAACGAAGTAACTTTTCCTTTCTCCTA
CCTAACTCAAAGATTAGGCCGAGCGCGATTTACTGATTACTAAGTATCTTGATAATTATG
TTTTTACAAGAGTAAAAAAAAGCGCAATAAAGGTTAATGTTTATCGTTTGGTACTATTGT
TTTTTTAGCTGAAAGTTTAGAACTCTAACCTTTCAGCTAAAAAAATTGTATCGACTTTTT
TTTGTAACAATTTTGTATTATACTGTAGACAAAGATTAAACAATCTTAAGTTTATCTATT
TTTTAGCTTTGTTTCTTTATTTGGTAAGAATAGATTTACAATGTGAAAAAAATAAGGTCT
CGCTATGCTCCTCTATCGCCGGGCTGGCAGCACTGATAACGAAGTTGTAGCAGCGATCGA
AGCCCTTCTTGTTTTTAGAAAGTCAATAAGGCCCAATAGCTACTAAAACCATGTGGTATT
GTTCCAACCAACGAGGTTGTAGGATAAAAAATAATTTTTAAAAACAATCCCTTTTAAAGC
AATTTAATTATTTTTCATTAATTACAAATCTTTCCTTCCTT
psbK ==> start
ATGATTCTTTTGTTTGCAAAACTTCCTGAAGCCTATGCGGCTTTTGACCCTATTGTAGAT
GTTCTTCCAGTTATTCCAGTGCTTTTTTTACTTTTAGCTTTTGTGTGGCAAGCTTCTGTA
AGTTTTCGTTAA
psbK ==> end
AAGTTTAGATTACGTAATTTAAATTACTCGTTTTCTTTAAAACATTCTAAATTTAATAAA
TTCATAGTATTTACATGAAGGAGTTTTAATTTGTTAAAGTGTTAAAAATTTAAGATGCTA
ACTTGTTTAAACAAAAGATTTTCAAGAATTTACAATACTAGAGGATTCTTTTTATTCGTT
ACTTGGCTACTACAAAAAAGTTTATAACAAAAATAATCTTTGTTGCAAATATGTAGTTAG
TCTTGTTACCAATCAACAGTTTTTGCAACTTAACCACTTGTTGAGAAACAACAAACGTTA
CTTTTGTTTGTGTTTTATATTGAGTGTTGATATGTATTGTAAATTTTATAAAATTCTTTT
ATTTTTCGGTTACTTAGTTCTTTTTTAATCAAACACTATTTTCCTTAAAAATTTCCCTTT
TTCTTGAGGGTTAAGATTTTATGGCT
psaM ==> start
ATTTCTGAAAGTCAAATTTTCGTAGCACTTTTTGCTGCTTTAATAACAGGTGTTCTTGCT
GTCCGACTTGGTATAGAACTCTATAGTCAACCTTAG
psaM ==> end
TTCTAGAAAATATAATATTGCGAATCATGTTCTTGATTCTTAAGAGTAACATAGAACGAT
TTGTTGATTTGAAAGCACATTTACAAATAAGCTAAATTTTTTCTCATTTTCTATACGATC
AAGCTTCAATAAAAAATTTGCATTAATTTTGCAATGTAAAGAAGCTTCACCTACATGCTT
TTGAGACTACTACATTTTTAATGTAGTAGTCCCAAAAAAAGATTCTGTTTCTTACAAAAT
TCAGATTTTAGTCTGAAGAATGTAAATCTTTTATACCTTGTCAACCCTCTAGGGTATAAA
GCAAATGAAACTCTTCTTTCAAGAAAATGTTAAATTTAGCAAAAACAGGTCTATGTAGCT
GTTTCTCTCTTATTCCATGCTTTTTAAATTTACAACAATAGGGCCTGCTTTCCTACCCAG
ACCCGTACCCTTTAGAGTTAGAAGCAGCCTTTCAGGTCTGGGTAGGAAAGCAGCGCTGAC
AGCGCCGGGCACATCACATGGAGATATATCAGCTAACTGTTTGTTTAGTAGTTGCCCTGA
TATAAATTTTCCGTTTGTCAGAGCACAACAAAGTAAAGAATTATACTTTTCTTAATATAA
ATACTTTTTGAAAATATAGTTGTAAAACATTGTAACTAATTGTAACATTAACTAGTAATT
TACTAAGAATTGTAGTATTA
trnL(gag) ==> start
GCCCTTGTGACGGAATGGTAGACGTGCCGTGTT!GAG!GTCGCGGTGCTTTTTAGTGTGT
CGGTTCGACTCCGACCAAGGGTA
trnL(gag) ==> end
CTTTCATTCAAATTCTTAGACTGTCATTGCTGTCATTGCTGTCAGCGCTGACAGCAAT
```
